# Scalable Plasmonic Metasurface-Enabled Physics-Guided Self-Supervised Cellular Imaging

**DOI:** 10.64898/2026.06.21.733589

**Authors:** Cheng Zhang, Susobhan Choudhury, Kerstin Jansen, Johannes Balkenhol, Katrin G. Heinze

## Abstract

High-quality cellular imaging, especially in live cells, remains constrained by the trade-off among signal-to-noise ratio, phototoxicity, and instrumentation complexity. Here, we report a scalable plasmonic metasurface that generates a spatially ordered array of fluorescence-enhancing near-field hotspots and enables self-supervised denoised, cellular imaging with improved feature readability on a conventional wide-field microscope. The registered hotspot lattice serves as a physics-derived functional prior that identifies where fluorescence amplification is physically grounded and steers neural-network training accordingly, reducing reliance on paired ground truth, large external pretrained models, or extensive supervised datasets. We demonstrate two labeling-density-dependent operating regimes: dense labeling for cytoskeleton structural imaging and sparse labeling for multiplexed sensing of plasma-membrane-associated dynamics across the hotspot array. Our work unites scalable nanophotonic hardware and self-supervised computational imaging into a practical platform for structural bioimaging and on-chip live-cell biosensing under simple wide-field imaging conditions.

## Introduction

Optical microscopy lays the foundation for modern cell biology, yet many molecular processes unfold at spatial scales beyond the reach of conventional diffraction-limited imaging. Although super-resolution fluorescence methods have greatly expanded the spatial information accessible to optical imaging, their broad adoption—especially for live-cell studies—remains limited by trade-offs among resolution, imaging speed, photon budget, phototoxicity, sample preparation, and instrumental complexity^1–6^. In practice, higher resolution often comes with a stronger experimental burden, either through complex optics, stringent labeling requirements, or demanding reconstruction pipelines^6–8^. Therefore, obtaining nanoscale biological information with conventional microscopes, minimal specimen perturbation, and practical experimental workflows remains a central challenge.

Plasmonic nanoantennas offer a distinct physical route to nanoscale optical interrogation^9^. This ability to localize and amplify light–matter interaction has made plasmonic nanostructures attractive for fluorescence imaging and molecular sensing at dimensions inaccessible to conventional far-field optics. Plasmonic structured illumination microscopy uses ordered nanoantenna arrays to generate sub-diffraction near-field excitation patterns that extend the spatial-frequency support of wide-field fluorescence imaging^10–14^. This capability depends critically on highly ordered and spatially uniform nanoantenna lattices, whose geometry and periodicity determine illumination fidelity and large-area imaging performance. In biosensing and single-molecule analysis, plasmonic nanoantennas, particularly nanoaperture-based platforms, have demonstrated that optical confinement can overcome the concentration and background limits of diffraction-limited fluorescence measurements. Therefore, they enable localized interrogation in environments ranging from biomolecular solutions to live-cell membranes^15–20^. These advances also reveal a central translational bottleneck. Many plasmonic applications depend critically on nanometer-scale pattern fidelity, while the fabrication methods that provide such precision—especially electron-beam lithography and focused-ion-beam milling—are typically slow, costly, and poorly suited to large-area, high-throughput manufacturing^21, 22^. As a result, the field is increasingly moving toward highly ordered and fabrication-tolerant plasmonic metasurfaces. Such platforms preserve optical quality while offering a more scalable way toward routine bioanalytical workflows. Beyond optical enhancements, ordered plasmonic hotspot arrays can also provide spatially registered physical information that is naturally compatible with computational image reconstruction.

Computational imaging has emerged as a complementary way to improve resolution and sensitivity without adding hardware complexity. Convolutional neural network (CNN)-based models now play a central role in microscopy because they efficiently learn image priors for denoising, deconvolution, and super-resolution^23, 24^. Yet many such pipelines still depend on supervised paired datasets that are expensive to generate and closely tied to specific acquisition conditions, which limits transferability across experiments. Transfer learning can reduce the need for extensive task-specific data but may introduce limited generalization when acquisition statistics or biological morphology deviate from the source domain^25^. This has driven increasing interest in physics-guided and self-supervised learning, where image-formation models or acquisition priors are embedded into the network to reduce sample complexity, stabilize reconstructions, and improve efficiency^26–29^. The next step for computational microscopy is therefore not merely better reconstruction quality, but greater physical fidelity and data efficiency.

In this article, we show that a scalable plasmonic metasurface can provide a physics-derived prior for a self-supervised CNN framework, enabling denoised fluorescence imaging with improved structural readability on a standard wide-field microscope. The ordered hotspot array serves not only to locally enhance fluorescence, but also to define a functional mask that constrains learning to regions of physically supported near-field signal. This converts near-field enhancement into a built-in supervisory prior, thereby reducing reliance on large-paired datasets or external pre-training. The platform is scalable in fabrication, free of specialized optics, and compatible with routine fluorescence labeling. We further demonstrate two complementary operating regimes: dense labeling for structural imaging reconstruction and sparse labeling for multiplexed live-cell membrane dynamics mapping. Overall, the approach integrates scalable nanophotonic hardware with data-efficient learning into a practical framework for cellular imaging, hotspot-based live-cell sensing, and future application-specific bioanalysis.

## Results

### Localized-plasmon-guided self-supervised CNN concept

To establish a physics-derived prior for fluorescence-based imaging reconstruction, we interface labeled cells with a planarized, ordered plasmonic nanotaper metasurface, in which the hotspot-bearing nanotaper apexes define the cell-facing plane (Fig. 1a). The nanotaper architecture is chosen to concentrate light energy toward the apex, where strong electromagnetic field enhancement is generated and induces the localized fluorescence hotspot^30^. Under wide-field illumination, conventional diffraction-limited fluorescence is emitted. In contrast, fluorophores located near the near field of the taper apexes experience strongly enhanced emission. This produces a spatially ordered set of fluorescence hotspots whose positions are fixed by the metasurface geometry and whose intensity origin is governed by plasmonic near-field enhancement. Their amplitudes are determined by local near-field coupling, fluorophore occupancy, and the biological labeling distribution. The plasmonic substrate therefore does more than increase local fluorescence detectability. It also converts the ordered hotspot array into a spatially registered experimental prior for defining a functional fluorescence mask, see Fig. 1b. We use this mask as a physics-derived prior in the CNN pipeline to constrain reconstruction toward spatially reliable regions of the image. The plasmonic substrate therefore serves a dual role: it amplifies the detectability of local fluorescence features and simultaneously supplies an experimentally encoded supervisory constraint for image restoration. Reconstruction is thus anchored by near-field physics instead of being driven solely by statistical image correlations.

**Fig. 1.**
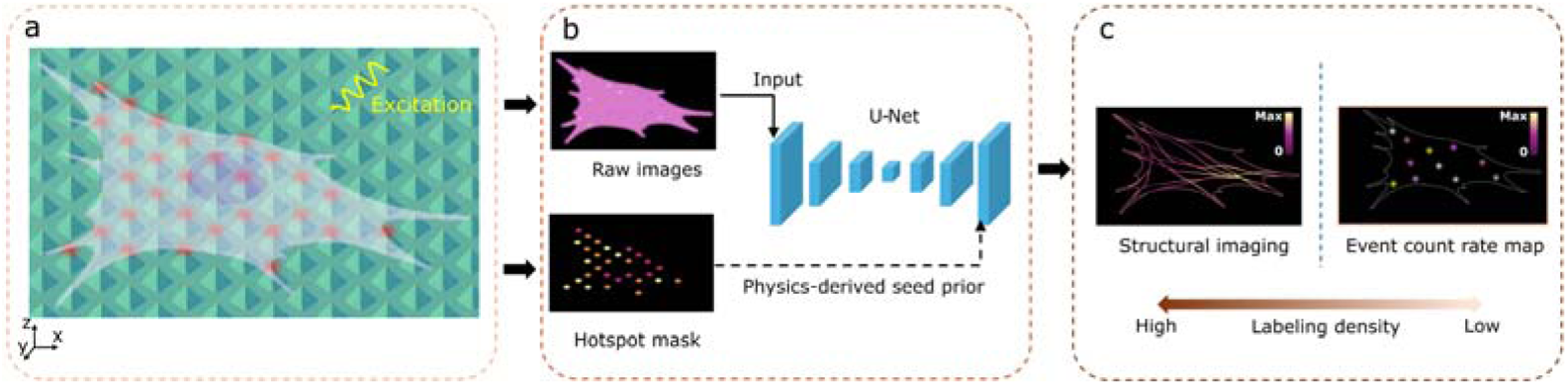
Concept of localized-plasmon-guided U-Net reconstruction for labeling-density-dependent cellular imaging. (**a)** A labeled cell is placed on a planarized ordered plasmonic nanotaper array, in which the nanotaper apexes define spatially predefined hotspot positions at the bio-interface. Upon wide-field excitation, fluorophores located near the plasmonic apexes generate locally enhanced fluorescence. The hotspot signals carry both fluorescence intensity and spatial coordinate information: their positions are determined by the ordered antenna lattice, while their amplitudes are modulated by localized-plasmonic near-field enhancement and fluorophore occupancy. (**b**) The hotspot-associated signals are registered to the fluorescence image and extracted as a hotspot functional mask. This mask serves as a physics-derived seed that identifies experimentally sampled, plasmon-enhanced locations and anchors the self-supervised U-Net reconstruction. In this way, the ordered plasmonic substrate provides hardware-encoded spatial priors that bridge the optical measurement and the computational reconstruction framework. **(c)** Labeling-density-dependent operation of the localized-plasmon-guided U-Net framework. Dense labeling populates many hotspots and supports denoised structural readout with improved morphology readability. Sparse labeling produces isolated hotspot events. This mode is less suited for complete structural recovery, but it supports localized plasma-membrane-dynamics mapping at selected hotspot nodes.

We employ U-Net as the central computational backbone because its encoder–decoder architecture is widely used for image denoising and segmentation in microscopy^23, 31^. In the present localized-plasmon-guided U-Net, reconstruction is not driven only by statistical similarity between images. It is also constrained by the experimentally measured hotspot prior. This links the optical measurement to the computational reconstruction. It also reduces the dependence on paired ground-truth images, large supervised datasets, or external pretrained models. The resulting framework can therefore be viewed as a plasmonic-hardware / U-Net-software platform, while the operative mechanism remains localized-plasmon-guided learning. The operating mode of the localized-plasmon-guided U-Net is defined by the imaging objective—either structural imaging or sparse hotspot-based membrane dynamics mapping—and the corresponding labeling density is chosen to support that objective (Fig. 1c). For structural imaging, high labeling density increases hotspot occupation across the ordered lattice, providing dense sampling of the cell-substrate interface and enabling reconstruction of extended cellular architecture. For sparse hotspot-based readout, lower labeling density yields isolated, high-contrast hotspot-associated signals that do not support complete whole-cell structural recovery but remain suitable for nanoscale mapping of plasma-membrane dynamics at selected hotspot locations. Overall, the established platform therefore supports two complementary regimes: a structure-reconstruction regime at high labeling density and a dynamic-sensing regime at low labeling density.

### Scalable plasmonic metasurface characterization and imaging setup

We fabricate a plasmonic substrate that is compatible with simple fluorescence imaging (see Fig. 2a). Imaging is performed on a conventional wide-field microscope using regular LED lamp illumination. The sample is assembled as a flow-chamber geometry with a capping coverslip, while the plasmonic substrate is mounted facing downward toward the cell and buffer compartment. In this configuration, fluorophores near the cell-substrate interface couple directly to the localized near field at the exposed nanotaper apexes. The ordered nanotaper metasurface is designed for scalable fabrication (Fig. 2b). We fabricate the array via a combination between nanosphere lithography^32^ and template stripping^33^ , as shown in Figure S1. This strategy enables high-throughput production of quasi-ordered hotspot arrays over macroscopic areas. Importantly, the cell-facing apex side is defined by template stripping, which preserves an ultrasmooth Au surface with roughness of 0.38 nm (see Figure S2). Representative images (see Fig. 2b) across multiple length scales show the structural hierarchy of the metasurface: a sharply defined single nanotaper structure, a laterally extended quasi-ordered array, and a macroscopic substrate footprint. This combination of local structural precision and large-area coverage is essential for translating plasmonic hotspot platforms from small-scale nanophotonic demonstrations to practical bioimaging substrates.

**Fig 2.**
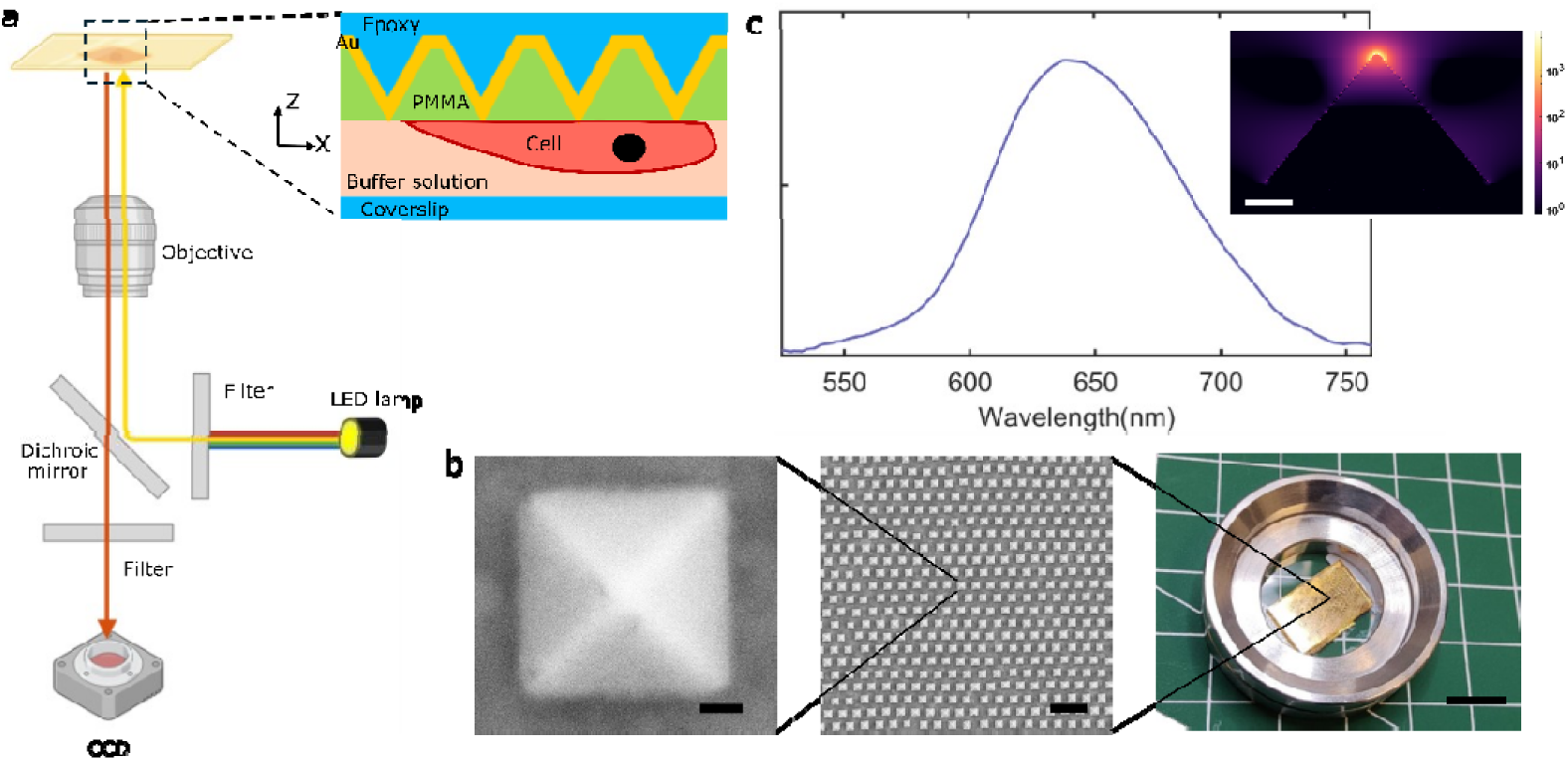
Scalable plasmonic nanotaper metasurface and optical characterization. **(a)** Experimental configuration for fluorescence imaging on a conventional wide-field microscope with LED excitation. Inset: cross-sectional sample geometry. The Au nanotaper array is planarized in PMMA and sealed with a capping coverslip to form a flow chamber. The plasmonic substrate is mounted upside down, so that the exposed nanotaper apexes face the cell-buffer interface. (**b)** Multiscale structural characterization of the nanotaper metasurface fabricated by nanosphere lithography and template stripping. Left: representative scanning electron microscope (SEM) image of a single nanotaper (pyramid bottom size of 240 nm). Middle: SEM image of a quasi-ordered nanotaper array over an extended area. Right: photograph of the macroscopic substrate, highlighting scalable large-area fabrication. Scale bars are 50 nm (left), 1µm (middle), and 1 cm (right), respectively. (**c)** Optical characterization of the nanotaper metasurface via white-light scattering measurement, showing a broad plasmonic resonance with a peak centered 640 nm. Inset, simulated local field-intensity distribution of the structure cross-section at 640 nm; the color bar is shown on a log scale. Scale bar, 50 nm.

The optical properties of the nanotaper metasurface are shown in Fig. 2c. The measured scattering spectrum exhibits a broad plasmonic resonance centered near 640 nm. Finite-difference time-domain simulation at the resonant wavelength shows a strong field localization at the nanotaper apex, where the local electric-field intensity is enhanced by more than three orders of magnitude with respect to the incident field. The combination of resonance matching, extreme field confinement, and scalable low-roughness fabrication establishes the nanotaper metasurface as the physical basis for robust localized plasmon guided fluorescence enhancement.

### Localized-plasmon-guided U-Net for the fixed-cell structural imaging

In this section, we demonstrate fixed-cell structural imaging in a dense-labeling regime on the nanotaper metasurface. Fixed CHO-K1 cells are cultured directly on the plasmonic substrate, labeled for F-actin with Atto 647, and imaged on a wide-field microscope at 639 nm using a low excitation power density of 1-2W/cm^2^. Under dense labeled conditions, a large fraction of nanotaper hotspots is expected to encounter nearby fluorophores^35^. This expectation is further supported by the occupancy analysis in Figure S3, which shows that for labeling densities on the order of 10^4^ molecules/μm^2^, the hotspot occupation probability approaches unity. Moreover, the mean fluorescence intensity averaged over the cell region is enhanced by approximately 3- to 5-fold relative to the control Au planar substrate without plasmonic nanotapers (see Figure 4S), in line with an earlier study showing plasmon-enhanced fluorescence imaging of labeled F-actin on nanostructured metallic interfaces^36^. The fixed-cell dense-labeling regime is therefore well suited for structural imaging. In this regime, the plasmonic lattice functions not only as a fluorescence-enhancing interface but also as a spatially defined sampling scaffold.

The raw fluorescence image already reveals the overall actin organization, but its interpretability remains limited by noisy background and by the spatially nonuniform coupling of fluorophores to the near-field hotspots (Fig. 3a). To introduce an experimental prior for reconstruction, we first co-register the hotspot lattice using the light scattering image of the underlying nanotaper array. Fluorescence-active hotspot sites are then identified and converted into an intensity-weighted functional hotspot mask (Fig. 3b and Figure S6). The hotspot mask initializes the learning as a seed target, and the reconstruction of the later iteration progressively expands the local support around those fixed seed positions, allowing continuous filament segments to be recovered beyond the original grid points (see supplementary note 8). In this manner, the ordered plasmonic lattice provides an externally encoded supervisory prior for the localized-plasmon-guided U-Net.

**Fig. 3.**
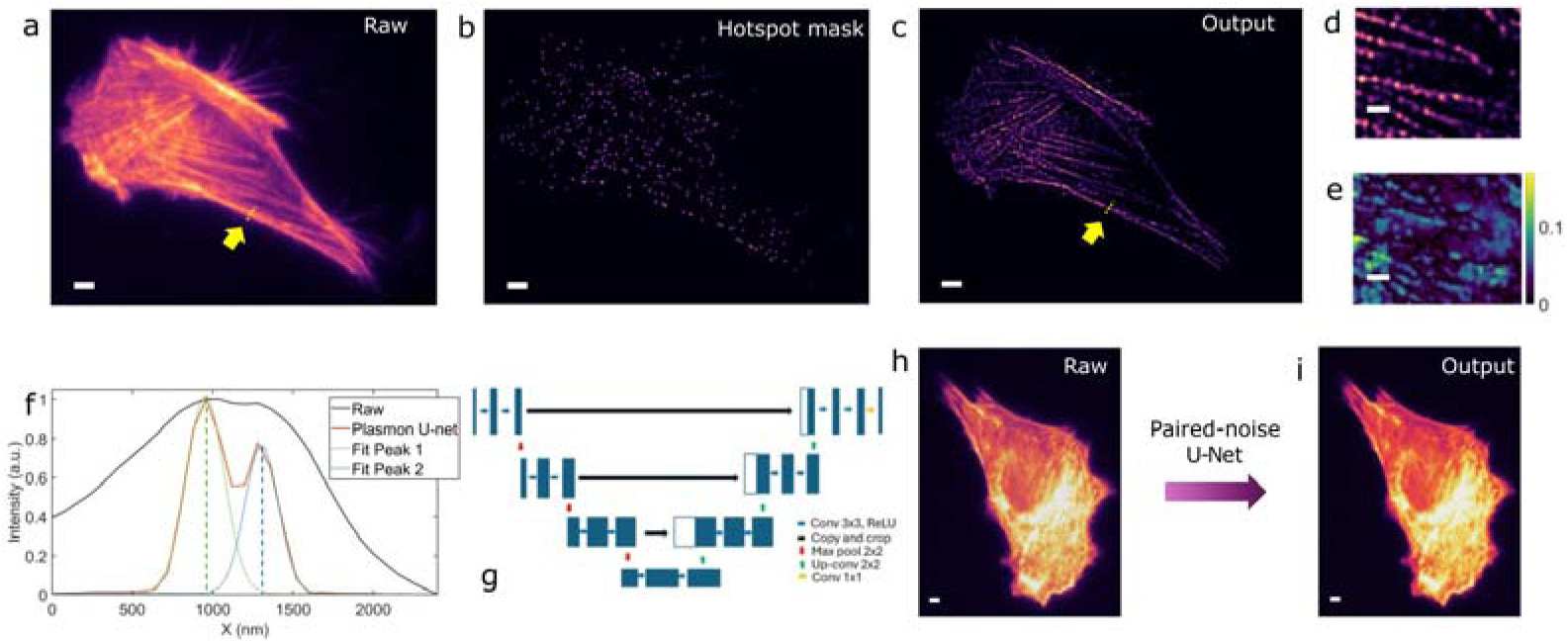
Localized-plasmon-guided U-Net for structural reconstruction in the fixed-cell dense-labeling regime. (a) Raw wide-field fluorescence image of an Atto 647 labeled F-actin fixed Chinese Hamster Ovary (CHO)-K1 cell acquired on the plasmonic metasurface substrate. Scale bar is 5 µm. (b) Intensity-weighted functional hotspot mask extracted from the co-registered scattering image and fluorescence response of the active hotspots. Scale bar is 5 µm. (c) Reconstructed output showing strong background suppression and enhanced recovery of filamentary actin structure. Scale bar is 5 µm. (d) A smaller area for showing more detailed reconstruction morphology. Scale bar is 2 µm. (e) Corresponding reconstruction error map based on the NanoJ-SQUIRREL algorithm^34^. The global RSP is 0.84 and the RSE (resolution scaled error) is 0.088. Scale bar is 2 µm. (f) The line-cut (indicated by the yellow arrow in raw and output images) comparison between the raw input and the reconstructed output, with a two-peak fit indicating improved feature separability following localized-plasmon-guided U-Net reconstruction. (g) Localized-plasmon-guided U-Net architecture used for reconstruction based on a three-level encoder-decoder design with repeated 3 × 3 convolution + Rectified Linear Unit (ReLU) blocks, 2 × 2 max-pooling, 2 × 2 up-convolution, and a final 1 × 1 output layer. (h-i) Control images acquired without plasmonic guidance and processed using a paired-noise U-Net model, showing comparatively limited denoising and structural recovery in the absence of a plasmonic physical prior. Scale bars are 5 µm. As a benchmark, the reconstructed output has a global RSP of 0.95 and RSE of 0.13. Please note that a higher RSP does not necessarily imply a more physically faithful reconstruction, because methods that remain closer to the raw diffraction-limited input can score highly while showing less improvement in local feature separability.

The resulting reconstruction yields a substantially cleaner representation of the actin cytoskeleton (Fig. 3c,d). Relative to the raw image, the localized-plasmon guided U-Net output exhibits pronounced suppression of diffuse background together with preserved continuous filament trajectories across the cell. Quantitatively, the reconstruction retains strong agreement with the input, yielding a global resolution-scaled Pearson (RSP) coefficient of 0.84, consistent with high structural fidelity after localized-plasmon-guided restoration. From Fig. 3e, the residual errors are localized and patchy, indicating the main filamentous features are preserved. The network therefore does not simply increase global contrast. Instead, it preferentially enhances line-like features consistent with filament morphology while attenuating weakly structured haze. This behavior is physically consistent with near-field sampling. Plasmonic field enhancement decays rapidly with increasing distance from the substrate. The reconstruction is therefore expected to emphasize basal actin segments that lie closer to the hotspot plane, whereas structures farther from the interface contribute less strongly. In this sense, the ordered plasmonic substrate transforms the problem from unconstrained denoising into physics-guided structural readability enhancement. This interpretation is further supported by the line-cut analysis (Fig. 3f). The line-cut indicates improved local feature separability after localized-plasmon-guided U-Net reconstruction, with representative filament-associated FWHM around 250 nm and an adjacent-feature separation of 350 nm becoming clearly distinguishable. Additionally, the importance of correct physical registration is tested by replacing the registered hotspot prior with a randomly shuffled, nonphysical mask. Under this condition, the reconstruction deteriorates markedly and produces flattened, spurious features (Figure S7). Therefore, successful recovery depends on correct hotspot registration, not on arbitrary masking. Although no paired high-resolution ground truth is available, the reconstruction is conservatively anchored to experimentally registered fluorescence-active hotspots. The method is also data efficient. When the training raw stack is reduced from 1000 frames to 100 frames, while keeping the number of training epochs unchanged, the reconstruction remained qualitatively similar with only a minor loss of sharpness (see Figure S8). Furthermore, another independent fixed-cell example shows comparable structural readability enhancement (see Figure S9), supporting the robustness of the workflow.

As a benchmark, we also trained a conventional self-supervised paired-noise U-Net, inspired by the Noise2Noise model^37^, using even/odd noisy fluorescence frames as input and target on a planar Au control substrate(see Fig. 3h-i). The model produces only modest denoising, mainly suppressing diffuse background noise while showing limited recovery of fine cellular morphology. Such behavior is expected because a paired-noise model relies only on statistical consistency between noisy observations and lacks an external physical prior that specifies where reliable signal should be preserved.

In our fluorescence stacks, temporal fluctuations and residual inconsistency between frame pairs further favor conservative global smoothing. This same mild global smoothing behavior was also observed when the paired-noise U-Net was applied to fluorescence stacks acquired on the plasmonic substrate itself, indicating that the limited recovery mainly reflects the unguided paired-noise framework rather than the substrate condition alone. Collectively, these results establish the dense-labeling fixed-cell regime as a principal operating mode of the localized-plasmon-guided U-Net, in which high hotspot occupancy enables broad near-field sampling of the basal cytoskeleton and links nanoscale optical enhancement to image-domain structural reconstruction.

### Localized-plasmon-guided U-Net for live-cell multiplexed membrane dynamics mapping

$Fig. 4 presents the live-cell sparse-readout mode of the localized-plasmon-guided U-Net, in which the ordered plasmonic lattice operates as a distributed nanoscale grid for membrane dynamics mapping. In this regime, the biological objective is not structural reconstruction but spatially multiplexed readout of localized membrane-associated dynamics across the cell footprint. Live CHO-K1 cells are labeled sparsely for β2-adrenergic receptor using SNAP-tag 596, with an estimated labeling density of approximately 100 molecules /μm^2^. At this density, the expected occupancy of individual plasmonic hotspots remains low (e.g., <10%), favoring isolated hotspot-linked fluorescence events (see Figure. S3). Relative to the control substrate without plasmonic metasurface, the plasmonic substrate yields an approximately one-order-of-magnitude enhancement in the mean fluorescence signal within the cell region (see Figure. S11). Because of the sparse labeling condition, the excitation intensity is increased to the ∼10 W/cm^2^ (596 nm excitation) range to improve signal-to-noise ratio while maintaining an acceptable balance between SNR and live-cell stability.

**Figure 4.**
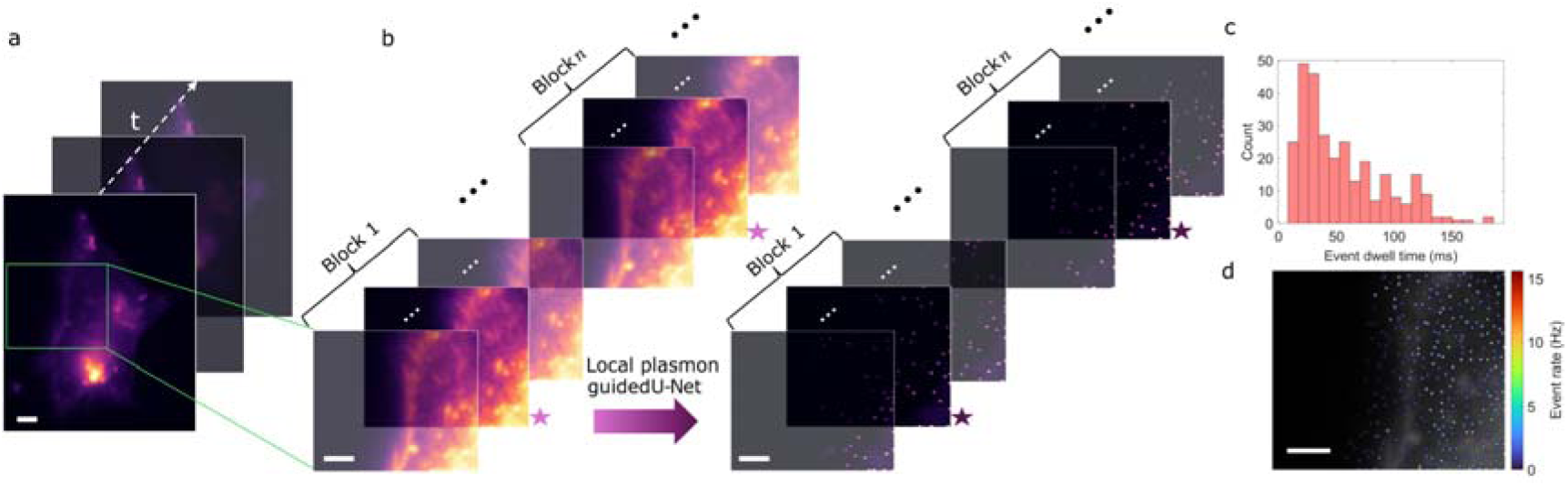
Live-cell localized-plasmon-guided U-Net for multiplexed plasma-membrane dynamics mapping. **(a)** Representative live-cell fluorescence time series acquired on the plasmonic metasurface under sparse SNAP-tag labeling of a β2-adrenergic receptor. **(b)** Principle of the block-calibrated live-cell sensing workflow. A selected cell region is divided into sequential temporal blocks; for each block, the central calibration anchor frame is used to locally calibrate the localized-plasmon-guided U-Net, and the calibrated model is then applied to generate a denoised sensing readout time series for that block. Here for clarity, block 1 and block n (n = 100, and it is not the last block) are shown here as illustrative examples. The central calibrated frame of the raw data is marked with light pink star while the output calibrated frame is marked with dark purple star. All the rest frames are set with a certain transparency only for visualization purposes. **(c)** Histogram of detected local plasma membrane dynamics event detection durations based on a one-minute movie. Events were defined from registered hotspot ROI traces after local baseline subtraction; frames with positive residual intensity and local signal-to-noise ratio (SNR) > 2 were classified as ON. Consecutive ON frames at the same hotspot were grouped as one event, and the event duration was calculated as the number of ON frames multiplied by the 10 ms frame integration time. **(d)** Event-rate map reconstructed at the functional plasmonic lattice sites, revealing spatially heterogeneous plasma-membrane dynamics across the cell footprint. An overlaid raw cell image is added as an easy guideline.

Representative raw live-cell fluorescence frames are shown in Fig. 4a. The corresponding workflow with a zoomed area is displayed in Fig. 4b. The time series is partitioned into sequential short temporal blocks (here, each block comprises 15 frames with each frame using 10 ms integration), and within each block the central calibration frame is used as a local anchor for block-wise calibration of the localized-plasmon-guided U-Net. The calibrated model is then applied to the frames of the same block to generate a denoised output. This strategy maintains local adaptation to evolving live-cell signal conditions while preserving the spatial registration imposed by the plasmonic hotspot lattice. In this sparse mode, the network functions primarily as a physics-guided activity extractor rather than as a generic denoiser: the ordered metasurface defines a large set of functional sensing nodes across the field of view, and the localized-plasmon-guided U-Net enhances the detectability of hotspot-supported membrane-associated events under photon-limited live-cell conditions (see supplementary note 12). The platform thereby enables parallel multiplexed readout across the cell footprint using a conventional wide-field microscope, without scanning instrumentation or externally supervised training data.

The detected event-duration distribution is summarized in Fig. 4c. A substantial fraction of events lies in the 20-50 ms range, indicating sensitivity to short-lived nanoscale membrane-associated dynamics. As each frame integrates for 10 ms and a single functional plasmonic hotspot probes an effective membrane area on the order of 40 nm in diameter, fast unconfined lateral diffusion of labeled receptors at sub-10 ms level through the hotspot is not expected to be fully resolved. The observed events are therefore operational hotspot-supported membrane-dynamics readouts, compatible with localized slowing, transient retention, repeated occupancy, or heterogeneous membrane organization^38^. Recent live-cell single molecule imaging reported sub-100 nm membrane compartments with a characteristic dwell time of sub-50 ms^39^, supporting that dynamics in this temporal range can arise in heterogeneous membrane environments. Fig. 4d converts the denoised hotspot detections into an event-detection-rate map across the functional plasmonic lattice. The resulting map reveals spatial heterogeneity in hotspot-supported membrane events across the cell footprint and illustrates the ability of the platform to provide simultaneous localized readout at many nanoantenna sites in parallel. In summary, these results establish the live-cell sparse-readout mode of localized-plasmon-guided U-Net as a practical framework for multiplexed plasma-membrane dynamics mapping that combines scalable nanophotonic hardware, conventional wide-field microscopy, and physics-guided learning.

## Discussion and conclusion

Here we demonstrate that the integration of scalable plasmonic nanophotonic hardware with physics-guided self-supervised learning can substantially extend the capabilities of conventional fluorescence microscopy. Importantly, the localized-plasmon-guided U-Net should be viewed primarily as a physics-constrained image-recovery framework, rather than a purely data-driven denoising approach. Unlike conventional deep-learning methods that rely on large, annotated datasets and may hallucinate structures absent from the underlying measurement, our approach incorporates experimentally defined optical information through the plasmonic hotspot lattice. Consequently, image reconstruction is guided by known physical constraints, improving robustness and interpretability while reducing dependence on extensive ground-truth training data.

In the dense-labeling structural-imaging mode, it is important to emphasize that the concept is not restricted to fixed-cell specimens. Fixed cells are intentionally selected here as a proof-of-concept validation system as they provide a stable and well-controlled platform for determining whether the plasmonic hotspot lattice can function as a reliable physics-derived prior for recovery of biological morphology. The successful reconstruction of features with improved local separability in this setting establishes the feasibility of the approach and provides a foundation for future applications in dynamic biological systems: Whenever labeling density is sufficiently high to populate the hotspot lattice, the same plasmon-guided reconstruction strategy can be applied.

At the same time, the current implementation has clear boundaries. Under the present optical configuration, the observed improvement is best described as enhanced structural readability and local feature separability rather than diffraction-unlimited imaging; it does not constitute a demonstration of diffraction-unlimited far-field imaging. This distinction is important because the hotspot prior is intentionally broadened through a Gaussian kernel to remain compatible with the optical characteristics of the water-immersion imaging systems. Consequently, the method improves morphology-oriented image recovery while maintaining a simple optical architecture and a low excitation burden. Nevertheless, several routes for further performance gains are evident. Improved hotspot definition, higher-numerical-aperture collection optics, greater hotspot uniformity, and tighter control of fluorophore–hotspot interactions should increase local feature separability. More broadly, the present work suggests a path toward hardware-assisted computational imaging in which engineered plasmonic excitation patterns encode spatial information directly into the measurement process. In future implementations, calibrated multi-pattern hotspot encoding may enable recovery of spatial information beyond the bandwidth accessible to conventional wide-field detection.

A second limitation concerns the sparse-labeling (live-cell) mode. Here, the network successfully extracts hotspot-supported events and enables multiplexed mapping of membrane dynamics across large fields of view. However, the reconstruction framework was optimized for morphology-oriented denoising rather than preservation of the full temporal statistics required for quantitative fluctuation biophysics. Neural networks that suppress noise and regularize frame-to-frame appearance can inadvertently alter local variance and temporal correlations, potentially biasing observables derived from autocorrelation or cross-correlation analyses^40^. This issue is particularly relevant under the present acquisition conditions, where molecular transit times may approach the camera sampling interval. Accordingly, the current implementation should be viewed primarily as a spatially multiplexed membrane dynamics sensing platform. Extending the method toward rigorous fluctuation biophysics will require temporally faithful network architectures, ideally incorporating fluctuation-aware forward models that explicitly preserve local variance, lag-dependent correlations, and causal timing, together with independent validation against raw-data observables.

In conclusion, we establish a scalable plasmonic metasurface as both a fluorescence-enhancing interface and an experimentally encoded physics prior for self-supervised microscopy. By coupling the ordered hotspot lattice with a localized-plasmon-guided U-Net, we demonstrate two complementary imaging modalities on a conventional wide-field microscope: morphology-oriented structural denoising in densely labeled samples and multiplexed membrane-dynamics sensing in sparsely labeled live cells. More broadly, this work highlights how nanophotonic hardware and physics-constrained artificial intelligence can be co-designed to extract information beyond that accessible through either component alone, providing a practical framework for future structural bioimaging and quantitative nanoscale bioanalysis.

## Associated content

### Data Availability Statement

The data supporting the key findings of this study will be deposited in Zenodo and made publicly available upon publication.

### Notes

The authors declare no competing financial interests.

### Author Contributions

C.Z. conceived the idea. C.Z. developed the plasmonic metasurface fabrication workflow and fabricated the samples. K.J. did the cell culture, fixation and staining for the fixed-cell imaging section. S.C. did the cell culture, membrane receptor labeling for the live-cell imaging section. C.Z. performed the experiment, analyzed data and built the U-Net working pipeline with assistance from J.B.. K. G. H. acquired the funding and supervised the project. All authors contributed to the discussions about this work, analysis of experimental data, comments on the manuscript, and writing of the manuscript.

## Acknowledgments

This work was supported by the Bavarian State Ministry of Science and the Arts supported K. G. H. (“Integrated Spin Systems for Quantum Sensors (IQ-Sense)”). Microscopy service was provided by the Core Unit Fluorescence Imaging in Rudolf Virchow Center for Integrative and Translational Bioimaging, Julius-Maximilians-Universität Würzburg (JMU). C.Z. thanks the financial support from the Rudolf Virchow Center for Integrative and Translational Bioimaging. C.Z. thanks Prof. Bert Hecht for accessing the white-light scattering measurement setup. C.Z. thanks Dr. Katherina Hemmen for the helpful comments on the manuscript. The authors thank the support from Wilhelm-Conrad-Röntgen-Forschungszentrum für komplexe Materialsysteme (RCCM), JMU for accessing the SEM imaging facility. The authors thank Prof. Jens Pflaum and Helena Hollstein for the initial helpful discussion.

## Notes

### Competing Interest Statement

The authors have declared no competing interest.

