## supplemental notes and figures for "Scalable Plasmonic Metasurface-Enabled Physics-Guided Self-Supervised Cellular Imaging"

**Supplementary notes:**

1. **Cell preparation protocols**
2. **Cell epi-fluorescence imaging setup**
3. **White light resonant nanostructure scattering measurement**
4. **Electromagnetic numerical simulation**
5. **Fabrication of plasmonic nanotaper metasurface**
6. **Labeling-density–dependent occupation of plasmonic hotspots**
7. **Fluorescence enhancement from fixed cell samples**
8. **Fixed-cell localized-plasmon-guided U-Net imaging pipeline**
9. **Odd/even paired-noise U-Net baseline model**
10. **Localized-plasmon U-Net image fidelity analysis**
11. **Additional information for fixed-cell data evaluations**
12. **Live-cell localized-plasmon-guided U-Net imaging pipeline**
13. **Computation hardware and software**

**Note 1: Cell preparation protocols**

1. **Fixed-cell preparation workflow**

CHO-K1 cells were cultured under standard conditions in the routine medium at 37 °C and 5% CO₂. Prior to cell seeding, plasmonic metasurface substrates, planar Au substrates, and glass coverslips were cleaned, rinsed with sterile Phosphate-buffered saline (PBS), and placed in a sterile culture dish. Cells were seeded onto each substrate at comparable density of 1×10^5^ cells/well in six-well plates and then cultured overnight in DMEM/F12 (PAN Biotech) supplemented with 10% (v/v) fetal calf serum (Biochrom), 100 U/mL penicillin, and 0.1 mg/mL streptomycin (Thermo Fisher Scientific) at 37 °C under 5% CO₂. Before cell fixation, the culture medium was then removed, and the samples were gently rinsed with warm PBS to remove non-adherent cells. Cells were fixed using freshly prepared 4% Paraformaldehyde (PFA, Thermo Fisher Scientific) solution for 15 min at room temperature. After fixation, the samples were washed three times with PBS. F-actin was stained using Atto 647N-phalloidin (Sigma-Aldrich) at a final concentration of approximately 10 nM in PBS-based labeling buffer for 30 min protected from light. After staining, the substrates were washed with PBS and stored in PBS at 4 °C sealed with aluminum foil until imaging.

2. **Live-cell preparation workflow**

A plasmid encoding an N-terminal SNAP-tagged β2-adrenergic receptor (SNAP–β2AR) was generated by inserting the SNAP-tag^1^ sequence upstream of the receptor coding region, as previously described^2^. CHO-K1 cells were cultured in DMEM/F12 (PAN Biotech) supplemented with 10% (v/v) fetal calf serum (Biochrom), 100 U/mL penicillin, and 0.1 mg/mL streptomycin (Thermo Fisher Scientific) at 37 °C under 5% CO₂. For transient transfection, cells were seeded at 1 × 10⁵ cells per well in six-well plates and incubated overnight. Transfection was performed using Lipofectamine 2000 (Life Technologies) according to the manufacturer’s protocol. Briefly, 250 ng plasmid DNA and 750 ng Lipofectamine 2000 were diluted in Opti-MEM (Thermo Fisher Scientific), combined, and added to phenol red–free DMEM/F12 to a final volume of 2 mL per well. Cells were incubated with the transfection mixture for 10–12 h. Following transfection, cells were washed, detached with trypsin, and reseeded onto plasmonic metasurface substrates, planar Au substrates coverslips and glass coverslips in phenol red–free DMEM/F12. After 4 h, coverslips were inverted and mounted onto HCl/NaOH-cleaned glass coverslips (24 × 40 mm) using double-sided tape spacers. Cell-surface SNAP–β2AR labeling was achieved using the cell-impermeable substrate SNAP-Surface 594 (New England Biolabs). Cells were incubated with 1 µM dye in complete medium for 20 min at 37 °C, followed by three washes with complete medium (5 min each). Imaging experiments were performed immediately after labeling.

**Note 2: Cell epi-fluorescence imaging setup**

Fixed and live labeled cells were imaged using a Nikon Ti2-E inverted microscope equipped with a standard epi-fluorescence imaging module. Excitation was provided by a multicolor LED source (Lumencor Spectra Light Engine, 8 channels), and fluorescence signals were collected on a scientific CMOS camera (C15440-20UP, Hamamatsu Photonics). Multiband Penta-bandpass filter sets were used for fluorescence detection, including spectral channels suitable for red to far-red regions. For fixed-cell measurements, samples were excited at 639 nm with an illumination intensity of approximately 1–2 W/cm² to limit bleaching and avoid excessive blinking-related fluctuations. Each captured frame used 10 ms integration. For live-cell measurements, 596 nm excitation was used at approximately 10 W/cm² to achieve a decent signal-to-noise during dynamic acquisition while maintaining acceptable live-cell stability. Each captured frame used 10 ms integration. Paired substrate scattering images were additionally recorded with integration time of 1 s, following the same detection pathway with fluorescence, in transmission geometry using white-light illumination from a halogen lamp.

**Note 3: White light resonant nanostructure scattering measurement**

We used a home-built white light scattering spectra measurement setup. A multimode fiber coupled from a halogen lamp is used as an excitation light source. The light beam is then focused into the back focal plane of an oil-immersion objective (Olympus 100×, NA = 1.45) to illuminate the sample with a collimated beam. A circular beam blocker is inserted in the detection path which allows for the collection of scattered light only. The scattered light is captured by a spectrometer (Shamrock 303i). In order to simulate the cell imaging environment, the samples were measured in an aqueous environment.

**Note 4: Electromagnetic numerical simulation**

Three-dimensional FDTD simulations were carried out using the Maxwell equation solver in the commercial software Ansys Lumerical. A high-resolution mesh (1 nm in all dimensions) was applied to the plasmonic nanotaper region to ensure accurate field resolution. Perfectly matched layers (PMLs) were implemented at all simulation boundaries to eliminate artificial back reflections. A plane wave is used to illuminate the nanotaper in the normal direction. An X-Z plan field monitor is used to visualize the field across the nanotaper. The dielectric function of the Au is based on reference^3^, the environment is water with refractive index of 1.3. The refractive index of epoxy substrate is set at 1.45.

**Note 5: Fabrication of plasmonic nanotaper meta-surface**

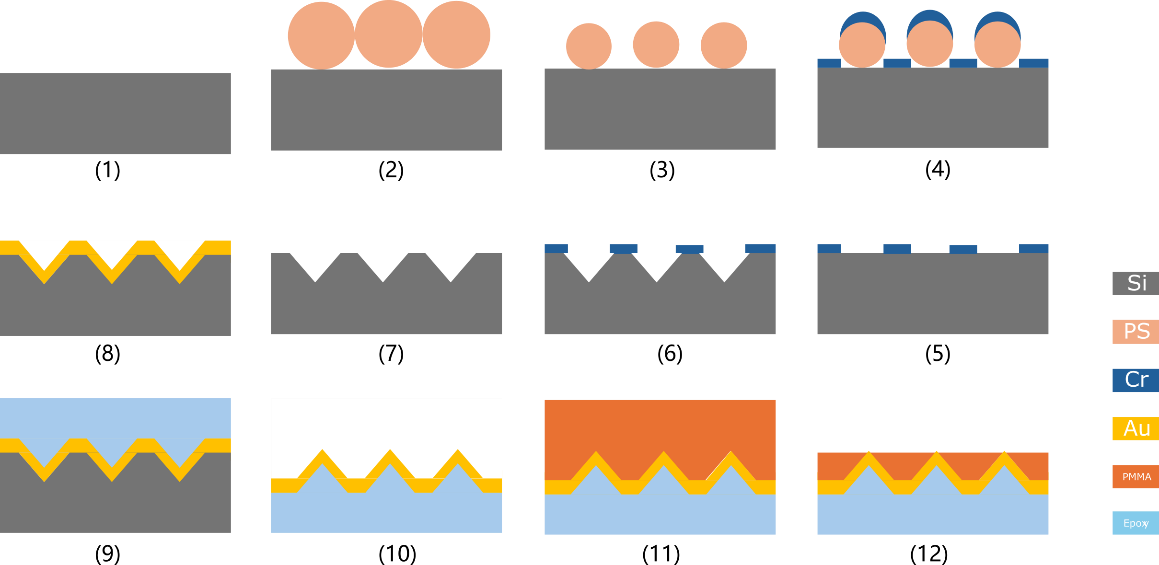

**Figure. S1 Nanofabrication workflow of the plasmonic nanotaper metasurface**

Plasmonic nanotaper metasurfaces were fabricated by combining colloidal self-assembly, anisotropic silicon (100) wet etching, metal replication, and template stripping. Silicon wafers were first used as substrates. A close-packed monolayer of polystyrene (PS) nanospheres was assembled on the wafer surface by Langmuir–Blodgett dip-coating (L-B Troughs, KSV NIMA, Biolin Scientific), providing a scalable colloidal mask over large areas. To reduce the sphere diameter and define the nanopattern mask geometry, the PS monolayer was subjected to oxygen plasma etching at 250 W, 10 sccm $O_{2}$, and 0.1 mbar for 12 min. A 20-nm chromium hard mask was then deposited by electron-beam evaporation at a base pressure of ${10}^{-6}$ mbar and a deposition rate of 1 Å/s. The PS nanospheres were subsequently removed in toluene, leaving behind a perforated Cr mask on the silicon surface. The pattern was transferred into the silicon substrate by anisotropic wet etching in 20 vol% KOH, yielding a periodic silicon nanotaper mold. After completion of the silicon etch, the remaining Cr mask was removed using a ceric ammonium nitrate-based chromium etchant. To form the plasmonic nanotaper structure, a 90-nm Au layer was deposited onto the silicon nanotaper mold by electron-beam evaporation at a deposition rate of 0.5 nm/s under a vacuum of ${10}^{-6}$ mbar. The Au nanotaper array was then transferred by epoxy-assisted template stripping. This step was used to expose the mold-defined Au surface, which provides substantially reduced roughness compared to the directly deposited top surface and is therefore favorable for plasmonic performance. To prepare the nanostructured substrate for biological experiments, the transferred Au nanotaper array was planarized with poly (methyl methacrylate) (PMMA) by spin-coating. Excess PMMA was then removed by oxygen plasma etch-back at 100 W, 10 sccm $O_{2}$, and 0.1 mbar for 6 min, stopping when the Au nanotaper tips became exposed. This procedure generated a quasi-planar surface in which the active Au plasmonic hotspots remained accessible, while the surrounding PMMA reduced topographical variation across the substrate. The resulting planarized nanotaper metasurface combined ordered plasmonic hotspots with a cell-compatible surface architecture and was used directly as a biocompatible substrate for cell culture and successive fluorescence imaging.

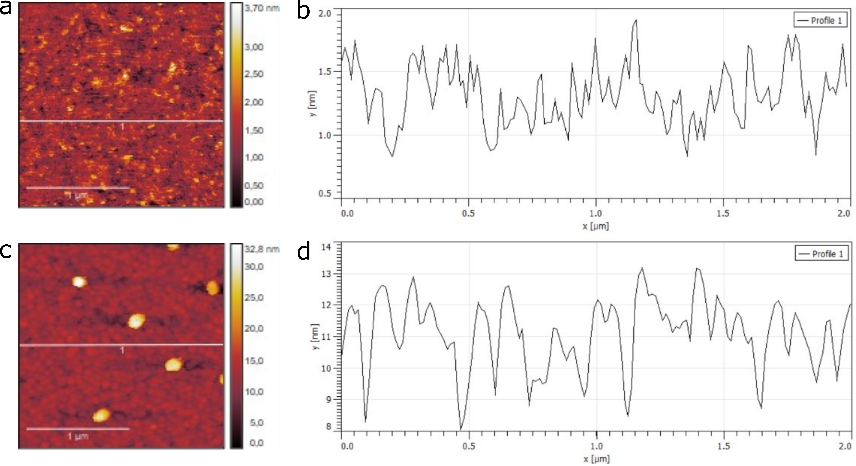

**Figure. S2** Atomic force microscope (AFM) characterization of surface quality between template stripped Au film from glass and direct evaporated Au film on glass. **a-b,** AFM surface image (2×2µm^2^) of the template stripped Au film and the corresponding line profile plot. **c-d,** AFM surface image (2×2µm^2^) of the directly evaporated Au film and the corresponding line profile plot. The mean roughness of the template stripped Au film is 0.38 nm, while the direct evaporated Au film has a roughness of 1.68nm.

**Note 6: Labeling-density–dependent occupation of plasmonic hotspots**

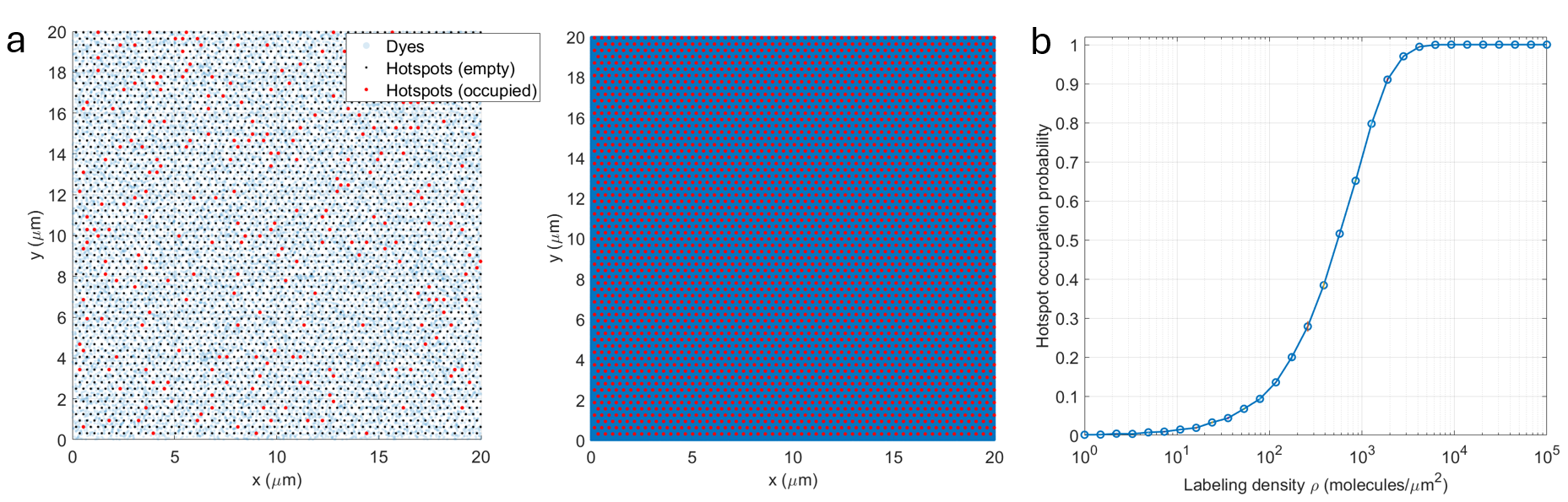

**Figure. S3. Labeling density governs plasmonic hotspot occupation and separates structural and sensing regimes**. **a,** Representative Monte Carlo realizations over a 20 × 20 µm^2^ region for low (left, 100 molecules/µm^2^) and high (right, 1×10^4^ molecules//µm^2^) labeling densities. Hotspots are arranged on a hexagonal lattice; fluorophores are randomly distributed. A hotspot is marked “occupied” if ≥1 fluorophore falls within the hotspot capture radius (effective near-field region), otherwise it is “empty.” At low density, only sparse hotspots contribute per frame; at high density, hotspots are nearly fully occupied. **b,** Hotspot occupation probability $P_{\mathrm{occ}}$versus labeling density $\rho$(log scale), computed as the fraction of occupied hotspots. Array disorder and axial dye–hotspot distance variations, which would reduce $P_{\mathrm{occ}}$, are intentionally neglected here for the sake of simplicity.

The fraction of plasmonic hotspots that contribute fluorescence in a given frame is primarily controlled by the labeling density of fluorophores. This provides a simple physical way to define two distinct operating regimes for our platform: (i) structural imaging under dense labeling, and (ii) sensing under sparse labeling. To quantify this transition, we model fluorophores as randomly and independently distributed on the membrane plane and treat each hotspot as “occupied” if at least one labeled molecule falls within an effective capture area $A_{\mathrm{eff}}$associated with the near-field enhancement region (Figure. S3).

Under these assumptions, the number of molecules within a hotspot follows Poisson statistics with mean $\lambda=\rho A_{\mathrm{eff}}$, where $\rho$is the labeling density (molecules/µm^2^). The occupation probability is then

$P_{\mathrm{occ}}=1-e^{-\rho A_{\mathrm{eff}}}.$	(S-0)

Using an idealized hotspot diameter of 40 nm (radius $r=20$nm) gives $A_{\mathrm{eff}}=\pi r^{2}\approx1.26\times{10}^{-3}{\mu m}^{2}$. This immediately yields the key scalings observed in Figure. S3: for sensing-level densities $\rho\lesssim100{\mu m}^{-2}$, $P_{\mathrm{occ}}\lesssim0.12$(e.g., $\rho=50{\mu m}^{-2}$ gives $P_{\mathrm{occ}}\approx0.061$), so only a small subset of hotspots is active per frame.

In contrast, for dense structural labeling typical of fixed-cell super-resolution^4^ (e.g., $\rho\sim{10}^{4}{\mu m}^{-2}$), $P_{\mathrm{occ}}\to1$. Consequently, essentially every hotspot is populated and emits with plasmonic enhancement, producing a quasi-continuous, hotspot-sampled representation of cellular structures. This is the regime where localized-plasmon-guided U-Net can most effectively leverage the ordered hotspot mask as a physics-derived prior to recovering structurally informative morphology over large fields of view.

Importantly, the curve in Figure. S3 represents an upper bound: in real samples, hotspot order is imperfect and the axial dye–metal separation varies across the membrane (linkers, membrane topography), both of which reduce the effective enhancement volume and thus decrease the apparent occupation probability. We neglect these effects here to isolate the first-order role of labeling density.

**Note 7: Fluorescence enhancement from fixed cell samples**

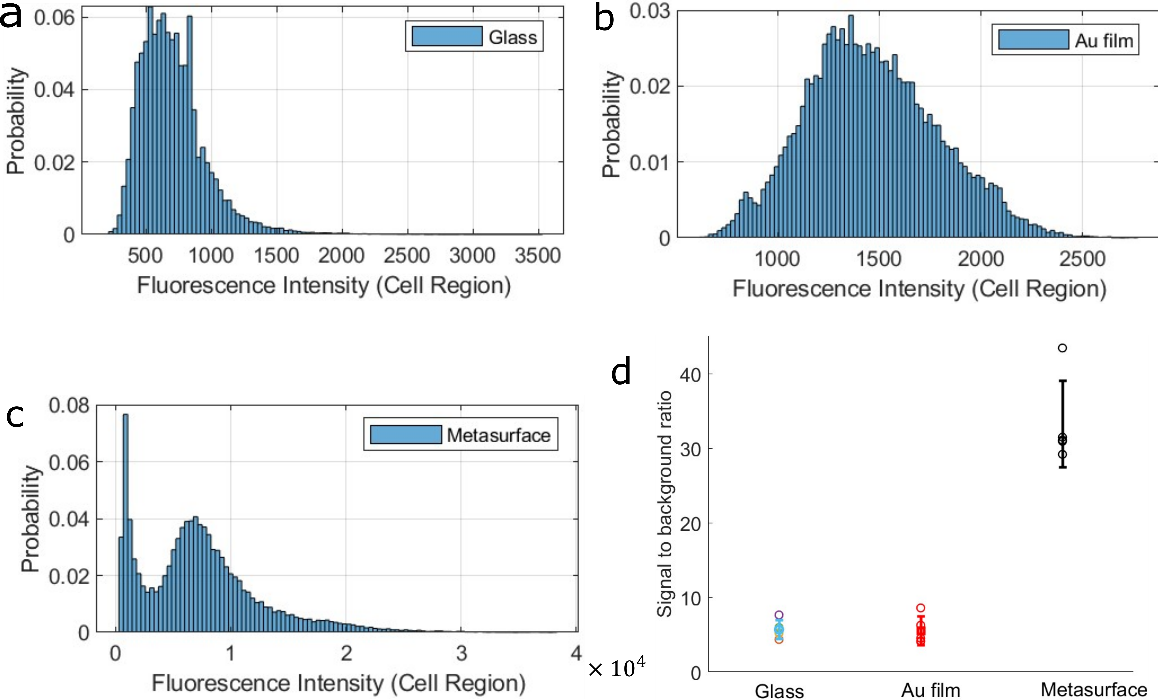

**Figure. S4. a–c,** Probability distributions of the cell-region-averaged fluorescence intensity measured from fixed cells on glass (a), planar Au film (with a 50 nm spacer PMMA layer) (b), and the plasmonic metasurface (c), acquired under matched imaging conditions. The glass control shows the weakest fluorescence distribution, largely below ~1000 counts, while the Au-film control exhibits only a modest shift to higher intensity. In contrast, the metasurface produces a strongly enhanced and broadened intensity distribution, extending up to ~10,000 counts. **d,** Statistical comparison of the signal-to-background ratio for the three substrates. The metasurface shows a clear and substantial increase over both control conditions. All fluorescence values were calculated as the average intensity within the segmented cell region.

To evaluate the fluorescence enhancement provided by the plasmonic metasurface under fixed-cell imaging conditions, we performed a comparative control study using three substrates measured under otherwise matched imaging settings: a conventional glass coverslip, a planar Au film, and the nanostructured metasurface. For each condition, the fluorescence signal was quantified by averaging the pixel intensity within the segmented cell region, thereby providing a substrate-dependent metric of the effective fluorescence output from labeled fixed cells. This analysis was designed to determine whether the observed signal gain originates simply from the presence of gold, or specifically from the nanostructured plasmonic architecture. The intensity distributions show a clear and systematic difference among the three substrates. Cells exhibit the smallest fluorescence intensity on glass, then on planar Au film with moderate increase, and most enhanced on the plasmonic metasurface substrate. This comparison is important because it separates the effect of material composition from the effect of nanoscale optical structuring. A planar Au film can modify the local optical environment, but it does not provide the strongly localized near-field confinement associated with plasmonic hotspots. By contrast, the metasurface concentrates the excitation field into spatially confined regions and can also alter the local emission process through enhanced light–matter coupling.

The statistical summary (Figure S4d) across all three conditions further confirms this trend. The signal-to-background ratio is lowest for glass and planar Au film, whereas it sharply increases for the metasurface. Such an increase is especially valuable for downstream image reconstruction and physics-guided learning, where robust signal contrast is essential for reliably identifying fluorescence events associated with the ordered hotspot array.

**Note 8: Fixed-cell structural imaging by localized-plasmon-guided U-Net pipeline**

For fixed-cell structural imaging, we employed a two-iteration physics-guided U-Net pipeline where the plasmonic hotspot map served as a seed prior. We used two experimental inputs: the wide-field raw fluorescence stack $X_{i}\left( r \right)$and light scattering image of the metasurface. Hotspot candidates were first extracted from the scattering image, filtered by the segmented cell region, and then matched to fluorescence local maxima in the fluorescence maximum projection. Only matched sites were retained as functional hotspots. Each hotspot was assigned a fluorescence-derived weight $w_{m}$, and the initial physics seed mask was constructed as a weighted Gaussian superposition

$M_{seed}\left( r \right)=w_{m}exp(-\frac{\left\| r-r_{m} \right\|^{2}}{2\sigma_{h}^{2}})$, (S-1)

where $r_{m}$denotes the functional hotspot coordinates, $\sigma_{h}=120nm$, and the whole functional seed mask is later normalized by min–max scaling. This Gaussian broadening was chosen to represent the optical hotspot footprint under the microscope with the water objective (NA=1.15). The first-iteration pseudo-target was then defined as:

$Y_{i}^{\left( 1 \right)}\left( r \right)=X_{i}\left( r \right)⨀ M_{seed}\left( r \right)$, (S-2)

with $\odot$denoting element-wise multiplication.

A U-Net with three encoder–decoder levels was trained to predict $\hat{Y}_{i}=f_{\theta}(X_{i})$ from the raw fluorescence input. $\theta$denotes the trainable parameters. Data augmentation during training consisted of random horizontal flips, vertical flips, and 90° rotations. Optimization was performed using Adam with learning rate ${10}^{-4}$and mini-batch size 15. In the present implementation, we optimized the global pixel-wise mean-squared error, thus, the loss function follows

$\mathcal{L=}\frac{1}{BN_{p}}\sum_{i=1}^{B} \sum_{p=1}^{N_{p}} \left( \hat{Y}_{i}(r)-Y_{i}(r) \right)^{2},$ (S-3)where $B$is the batch size and $N_{p}$is the number of pixels. The softly broadened plasmonic hotspot prior alone provided sufficient and more transferable guidance. By avoiding explicit structure-class priors, the reconstruction remained feature-agnostic while still allowing local biological continuity to emerge from the data.

To avoid over-constraining the first-pass output to the sparse initial seed, training was terminated using a soft hotspot-mask-weighted Pearson gate. After each epoch, the predicted validation maximum projection was Gaussian-degraded and compared with the raw maximum projection using a weighted pearson-like similarity, with weights derived only from the softly broadened hotspot mask,

$W\left( \mathbf{r} \right)=G_{\sigma_{m}}*M_{\mathrm{seed}}\left( \mathbf{r} \right),$ (S-4)where $G_{\sigma_{m}}$ is a mild Gaussian smoothing kernel. The corresponding gate value was computed as a weighted Pearson correlation between the raw projection and the degraded prediction under $W$, and a threshold of 0.95 was used as the empirical stopping criterion for the first iteration. Conceptually, this gate acts as an early safeguard: it limits excessive off-hotspot expansion before unsupported features can propagate into later refinement.

The first-pass prediction was then used to refine the support mask through controlled off-seed growth. After intensity matching the predicted maximum projection to the raw fluorescence projection, a growth term was extracted from the local structured contrast of the prediction, restricted to seed-connected regions and excluded from the seed itself,

$G_{\mathrm{grow}}(\mathbf{r})=\max(P(\mathbf{r})-(G_{\sigma_{\mathrm{bg}}}*P)(\mathbf{r}),\text{ }0)\cdot G_{\mathrm{conn}}(\mathbf{r})\cdot(1-M_{\mathrm{seed}}(\mathbf{r})),$ (S-5)where $P(\mathbf{r})$is the intensity-matched first-pass prediction, and $G_{\sigma_{\mathrm{bg}}}*P$provides a smoothed local background estimate, such that the positive contrast term extracts structured signal above diffuse haze. The seed-connected factor $G_{\mathrm{conn}}(\mathbf{r})$was obtained by morphological reconstruction using a dilated seed marker and the smoothed prediction as reconstruction mask, thereby retaining only structures connected to the hotspot-supported regions. Multiplication by $1-M_{\mathrm{seed}}(\mathbf{r})$further restricts the update to the off-seed region. As a result, $G_{\mathrm{grow}}(\mathbf{r})$ represents a controlled, seed-anchored off-hotspot extension term rather than unrestricted prediction-driven growth. The final mask was then defined as

$M_{\mathrm{final}}\left( \mathbf{r} \right)=\max\left( M_{\mathrm{seed}}\left( \mathbf{r} \right)+G_{\mathrm{grow}}\left( \mathbf{r} \right),\text{ }M_{\mathrm{seed}}\left( \mathbf{r} \right) \right)$. (S-6)

This formulation preserves the original hotspot prior as the highest-confidence anchor while allowing only controlled emergence of biologically meaningful off-seed structure.

To proceed, the refined support mask is used as the final training guide. The second-iteration pseudo-target was therefore defined as

$Y_{i}^{\left( 2 \right)}(\mathbf{r})=X_{i}(\mathbf{r})\odot M_{\mathrm{final}}(\mathbf{r}).$ (S-7)The network was then retrained using the same global mean square error loss. In this final stage, the mean loss decreased during the early epochs and then arrived at a stable plateau, indicating convergence of the refinement process (see Figure S5). The final fixed-cell output should therefore be interpreted as the result of an iterative physics-guided reconstruction in which experimentally registered plasmonic hotspots provide the transferable prior, while a conservatively updated refined support enables broader cellular continuity to emerge without introducing uncontrolled off-hotspot expansion in the final pass.

The final output should therefore be interpreted as the result of an iterative physics-guided feature-recovery process: experimentally registered plasmonic hotspots define the initial high-confidence seed prior, the first-pass network output expands this prior into biologically meaningful off-seed structures, and the refined guide then drives the final U-Net training toward a denoised, structure-preserving image of the fixed-cell architecture.

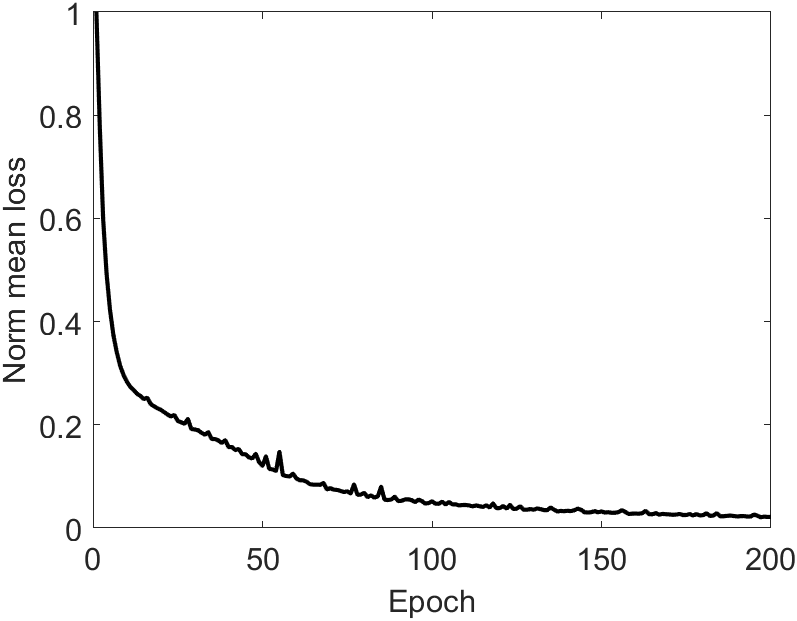

**Figure S5 | Mean loss evolution during final-stage training of the fixed-cell structural imaging pipeline.** Epoch-wise mean training loss for the final iteration of the physics-guided localized-plasmon-guided U-Net. After an initial decrease, the loss progressively approaches a stable plateau, indicating convergence of the network during refinement with the updated support mask. The observed stabilization is consistent with the final iteration acting primarily as a structure-consolidation step following seed-guided denoising and mask expansion in the first iteration.

**Note 9: Odd/even paired-noise U-Net baseline model**

As a generic self-supervised denoising reference, we implemented an odd/even paired-noise U-Net baseline. This model was intended to provide a full data-driven comparator without any physics-guided prior. The network followed a U-Net encoder-decoder architecture with skip connections, convolution, batch normalization, ReLU activation, max pooling in the encoder, transposed-convolution upsampling in the decoder, decoder dropout, and a final linear regression head. From the raw fluorescence stack, temporally adjacent odd/even frames were used to form noisy input-target pairs, from which spatially matched 96×96-pixel patches were randomly extracted and augmented by flips and${90}^{^{\circ}}$rotations. The model $f_{\theta}$was trained by minimizing the mean squared error

$\mathcal{L(}\theta)=\frac{1}{N}\sum_{i=1}^{N} \parallel f_{\theta}(x_{i})-y_{i}\parallel_{2}^{2},$ (S-8)where $x_{i}$and $y_{i}$denote paired noisy input and target patches, N is the number of training patch pairs. Training used the Adam optimizer with a minibatch size of 18 and an initial learning rate of ${10}^{-3}$. Validation loss on the held-out patch pairs was monitored during optimization. For full-frame inference, each image was denoised using overlapping tiled prediction to avoid memory limitations, and adjacent tile predictions were blended with a two-dimensional Hann window before stitching into the final output image. This odd/even paired-noise U-Net therefore served as a generic paired-observation self-supervised baseline, in contrast to localized-plasmon-guided U-Net, which additionally incorporates an experimentally measured plasmonic hotspot prior to guide restoration.

**Note 10: Localized-plasmon U-Net image fidelity analysis**

To assess reconstruction fidelity, we adapted NanoJ-SQUIRREL^5^ by comparing the diffraction-limited raw image with a resolution-scaled version of the localized-plasmon guided U-Net output. Following the NanoJ-SQUIRREL formalism, the reconstructed image was intensity-rescaled and convolved with a Gaussian resolution scaling function (RSF) to generate a diffraction-limited equivalent prior to comparison. Both raw and output images were first min–max normalized. The error map was computed as the pixel-wise absolute difference between the normalized raw image and the resolution-scaled output. Global similarity was quantified using the resolution-scaled error (RSE), defined as the root-mean-square difference, and the resolution-scaled Pearson coefficient (RSP), defined as the Pearson correlation between the normalized raw image and the resolution-scaled output. Because normalization was applied before evaluation, the reported RSE and error-map amplitudes are dimensionless and were used primarily for internal comparison across matched conditions, whereas RSP served as the main indicator of structural agreement. Moreover, because no paired higher-resolution reference image of the same cell is available, this analysis is used as an internal fidelity measure relative to the diffraction-limited input rather than as an absolute validation of structural truth.

**Note 11: Additional information for fixed-cell data evaluations**

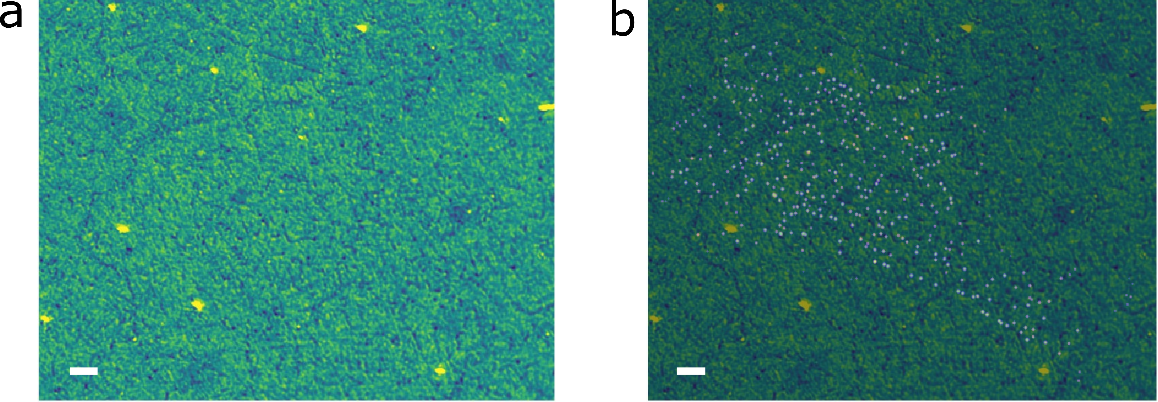

**Figure. S6** (a) The reflected scattering image of a quasi-nanotaper array; (b) overlay of the seed mask on the corresponding nanotaper scattering image, scale bar of 5 µm. Please note this underlying antenna array scattering image is only paired with its corresponding cell fluorescence image (in this case paired with the cell image in Fig. 3a).

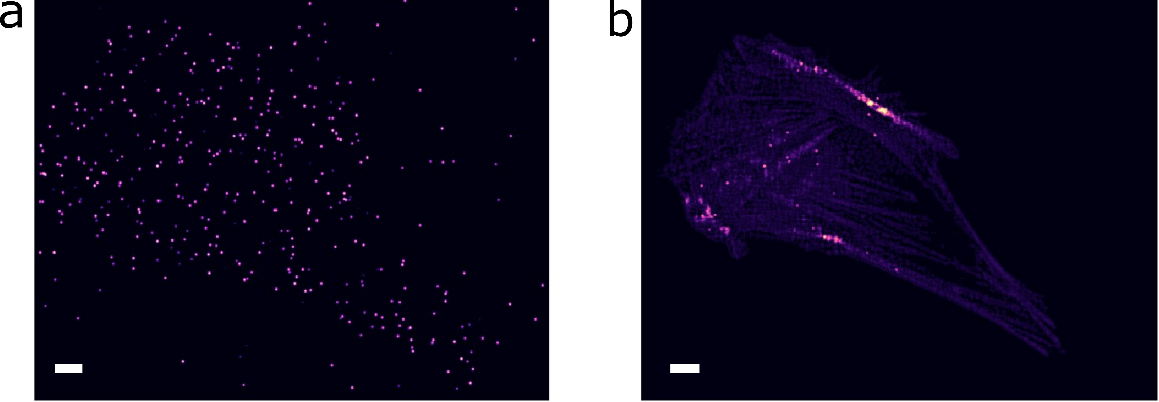

**Figure. S7** **Negative control of fixed-cell localized-plasmon U-Net reconstruction under a shuffled nonphysical hotspot mask**. (a) Shuffled hotspot mask generated by randomly redistributing hotspot coordinates across the full image plane (90% within the cell and 10% outside the cell). (b) Fixed-cell plasmon-U-Net output obtained using the shuffled mask as training prior. The reconstruction fails to preserve coherent filamentous architecture and instead shows flattened morphology together with numerous fragmented features. Scale bar of 5 µm.

To test whether a physically registered hotspot prior is mandatory for the fixed-cell localized-plasmon-guided U-Net, we performed a shuffled-mask control. The hotspot coordinates were randomly redistributed across the full image field before training. The shuffled mask was therefore no longer aligned with the registered plasmonic lattice. Under this nonphysical guideline, the reconstruction quality collapsed. Many filamentary structures appeared flattened or fragmented. Additional spurious punctate features also emerged, and fluorescence intensity across the cell is strongly mis-scaled. This stands in sharp contrast with the correctly guided plasmon U-Net output where extended actin filaments are preserved, and background noise is highly suppressed. This control therefore shows that successful reconstruction depends on correct physical hotspot registration, as opposed to arbitrary sparse masking or generic denoising.

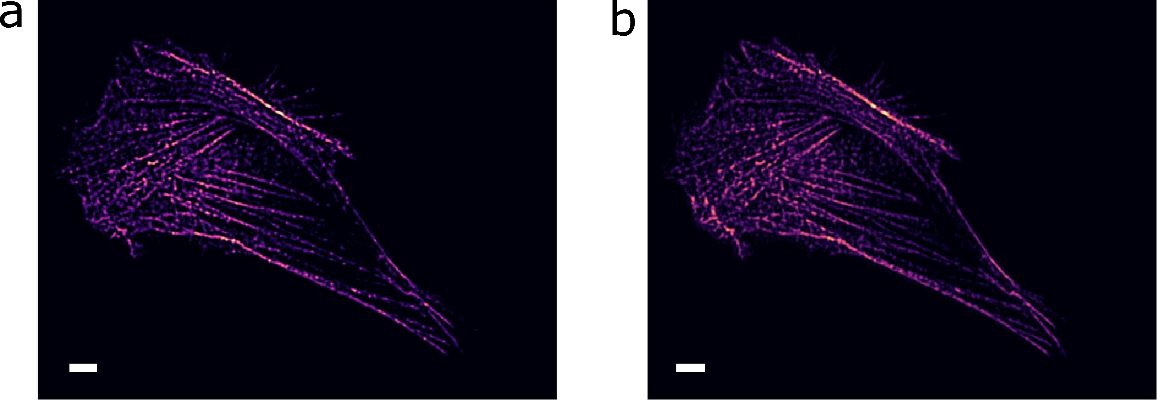

**Figure. S8 Validation of plasmon-U-Net performance under different training dataset volumes.**
Qualitative comparison of plasmon-U-Net reconstructions obtained after 200 training epochs using different numbers of training images. The model trained with 1000 images (left) and that trained with 100 images (right) both recover the major filamentous architecture of the cell with comparable overall morphology. Reducing the training set to 100 images leads to slightly softer biofeatures and a modest loss of the finest structural sharpness, but the main actin-like organization remains clearly preserved. These results indicate that the plasmon-guided U-Net is compatible with limited-data training, consistent with the stabilizing role of the hotspot physics prior. Scale bar of 5 µm.

To evaluate the dependence of localized-plasmon-guided U-Net on training dataset size, we compared reconstructions obtained after 200 epochs using either 1000 or 100 training images. The two outputs remain highly similar at the level of overall cell morphology and major filament organization, indicating that the network retains strong reconstruction capability even under markedly reduced data availability. When trained with only 100 images, the recovered biofeatures become slightly softer, with a modest loss of the finest structural contrast, but the principal actin architecture is still preserved. These results indicate that the plasmon-guided learning strategy is compatible with small-data training and benefits from the physics-derived prior encoded by the hotspot seed mask.

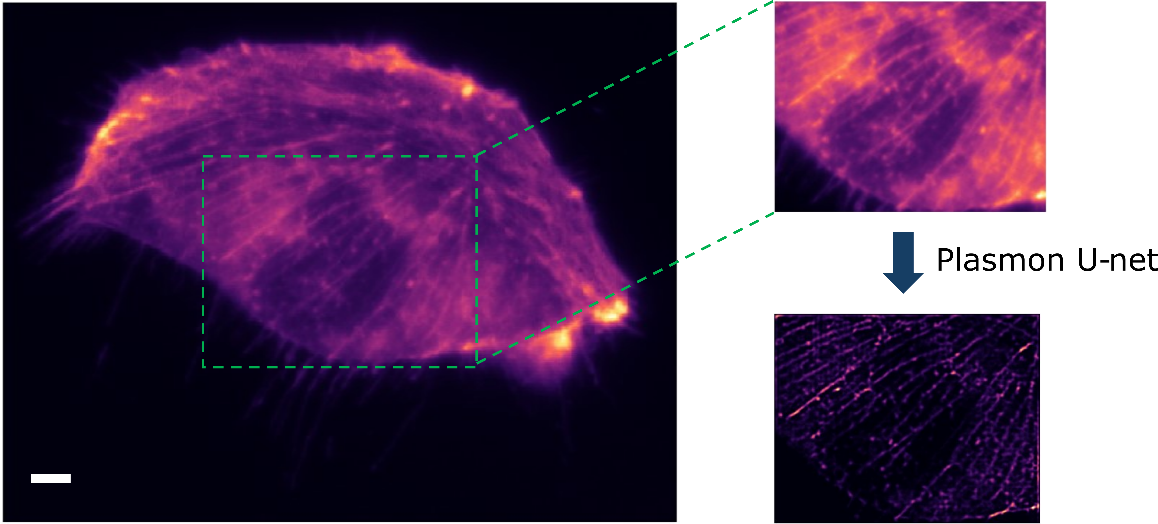

**Figure. S9** **Reproducibility of plasmon-U-Net reconstruction in an independent cell.** Representative reconstruction of a different cell processed using the same plasmon-U-Net workflow. The raw fluorescence image (left) exhibits diffuse background and limited structural contrast. A magnified region of interest (upper right) highlights the locally blurred filamentous features present in the input, whereas the corresponding plasmon-U-Net output (lower right) reveals stronger background suppression and improved recovery of the underlying filament network. The consistent denoising behavior observed in this additional sample supports reproducibility across cells with similar structural content of the plasmon-guided reconstruction strategy. Scale bar of 5 µm.

To assess reproducibility across biological samples, we applied the same localized-plasmon-guided U-Net workflow to an independent cell. The reconstruction again shows pronounced background suppression and improved delineation of the filamentous architecture relative to the raw fluorescence image. In the magnified region, weak and partially buried structural features become more clearly resolved after reconstruction, supporting the reproducibility of the denoising and feature-recovery behavior across different cells. This result suggests that the plasmon-guided framework is not limited to a single example but generalizes across samples with similar structural content.

**Note 12: Live-cell localized-plasmon-guided U-Net working pipeline**

For live-cell sparse sensing, we employed a block-wise calibrated localized-plasmon-guided U-Net in which the ordered plasmonic hotspot lattice served as a functional sensing prior rather than a structural-reconstruction guide. In contrast to the fixed-cell structural imaging pipeline, the live-cell implementation was not designed to recover the full cell morphology. Instead, it was designed to extract trusted hotspot-associated membrane dynamics from photon-limited fluorescence movies by restricting learning to hotspot-supported regions only. Consistent with the current executable implementation, only one learning iteration was used, so the live-cell model acts as a single-pass hotspot calibration and sensing readout.

The experimental inputs consisted of the wide-field raw fluorescence movie $R_{t}(\mathbf{r})$, an optional fluorescence movie used only for mask generation, and a white-light scattering image of the plasmonic metasurface. The scattering image was used to extract the static hotspot coordinates which define the physically registered sensing lattice. The fluorescence movie (1 minute) was then divided into temporal blocks of 15 frames. At 10 ms exposure per frame, each block therefore spanned 150 ms and was centered on a single calibration anchor frame. Within each block, the neighboring frames were not treated as equivalent training targets. Instead, they were used to construct a block-consistent cell mask, background template, and persistence support for hotspot validation and rejection of broad or cluster-like contamination. In this context, calibration refers to block-specific optimization of the network weights using the central calibration anchor frame, whereas the remaining frames in the block were processed only by inference under the fixed calibrated model.

For the calibration anchor frame, accepted fluorescence maxima were matched to the registered plasmonic hotspots and converted into an amplitude-weighted hotspot mask $M_{c}\left( \mathbf{r} \right)$followed by min-max normalization. In the present sparse-sensing implementation, the positive training support was restricted to the hotspot cores only. Thus, unlike the fixed-cell mode, no global morphological recovery term was intentionally introduced into the live-cell learning target.

We applied a synthetic cell region intensity degradation stack $\left\{ I_{k}(\mathbf{r}){\}}_{k=1}^{N} \right.$ from the central calibration frame $I_{c}(\mathbf{r})$only, with $N=200$ is the number of the total degraded frame number. For each realization, the network input was defined as

$\mathbf{X}_{k}(\mathbf{r})=\left[ \begin{aligned} I_{k}(\mathbf{r}) \\ M_{c}(\mathbf{r}) \end{aligned} \right],$	(S-9)

where $I_{k}(\mathbf{r})$ is the synthetic fluorescence realization. The corresponding pseudo-target was

$Y_{k}(\mathbf{r})=I_{k}(\mathbf{r})\odot M_{c}(\mathbf{r}),$	(S-10)

with $\odot$denoting element-wise multiplication. The U-Net prediction $\hat{Y}_{k}=f_{\theta_{b}}(\mathbf{X}_{k})$was then optimized using a hotspot-weighted masked mean-squared error,

$\mathcal{L}_{\mathrm{hot}}=\frac{\sum_{\mathbf{r}} W_{c}(\mathbf{r})\left[ \hat{Y}_{k}(\mathbf{r})-Y_{k}(\mathbf{r}) \right]^{2}}{\sum_{\mathbf{r}} W_{c}(\mathbf{r})+\varepsilon},$	(S-11)

where $\varepsilon$ is a small term to prevent division by zero; $W_{c}\left( \mathbf{r} \right)=\left[ G_{\sigma}*M_{c}(\mathbf{r}) \right]$ with$G_{\sigma}$ is a mild Gaussian smoothing kernel which represents softly broadened hotspot support derived from the accepted calibration-frame hotspots. In the current live-cell implementation, this hotspot-weighted masked loss was the only active optimization term, so that the training remained explicitly restricted to local hotspot-supported sensing readout.

Optimization was performed using Adam with learning rate ${10}^{-4}$, mini-batch size 5. Training was terminated once the hotspot-weighted masked mean-squared error showed no further substantial decrease and reached a stable plateau, a practical stopping point was typically reached at around 30 epochs. In representative runs (see Figure. S10), a rapid initial mean-loss drop within 5 epochs and thereafter a stabilized mean-loss value.

After calibration, the learned block-specific parameters $\theta_{b}$were held fixed and applied frame by frame to all frames within the same block. For each frame $t$, inference used the current raw image $R_{t}(\mathbf{r})$together with that frame’s own accepted hotspot mask $M_{t}(\mathbf{r})$,

$\mathbf{X}_{t}^{\mathrm{infer}}(\mathbf{r})=\left[ \begin{aligned} R_{t}(\mathbf{r}) \\ M_{t}(\mathbf{r}) \end{aligned} \right],\hat{Y}_{t}=f_{\theta_{b}}\left( \mathbf{X}_{t}^{\mathrm{infer}} \right).$ (S-12)Thus, the network weights were calibrated once per block, whereas the sensing mask remained frame-adaptive within the block.

A 15-frame block was used as a compromise between temporal fidelity and computational efficiency. 10 ms exposure per frame corresponds to a total duration of 150 ms and $\pm70$ms around the calibration anchor frame. Assuming a representative membrane receptor lateral diffusion coefficient^2^ $D=0.1\text{ }\mu m^{2}/s$, the two-dimensional root-mean-square displacement from the anchor frame to the block edge is about 0.17 $\mu m$ which remains below the $\sim0.4\text{ }\mu m$ nearest-neighbor hotspot spacing. The selected block length therefore provides sufficient temporal support for robust hotspot validation while remaining compatible with locally evolving receptor dynamics at the scale of the plasmonic lattice. In general, shorter blocks would increase the number of recalibration steps and the overall computation time, whereas longer blocks would increase the risk of losing rapidly varying hotspot-associated dynamics.

The final live-cell output should therefore be interpreted as the result of a block-wise calibrated physics-guided sensing process: experimentally registered plasmonic hotspots define the high-confidence sensing prior, the central calibration anchor frame determines the block-specific network weights through self-supervised hotspot-restricted learning, and the calibrated model is then applied to the remaining frames in the same block to generate conservative, denoised hotspot-event readout. For event analysis, fluorescence traces were extracted from fixed ROIs centered at the registered plasmonic hotspot coordinates, using the same hotspot geometry for raw and U-Net-processed movies. For each hotspot trace, a local baseline was estimated by a 1 s moving median, and local noise was estimated from the moving median absolute deviation of the baseline-subtracted residual; a frame was classified as ON when the residual was positive and exceeded the local noise by more than two-fold, corresponding to SNR > 2. Consecutive ON frames at the same hotspot were grouped as one operational hotspot-supported event, with the event duration given by the number of ON frames multiplied by the 10 ms frame integration time.

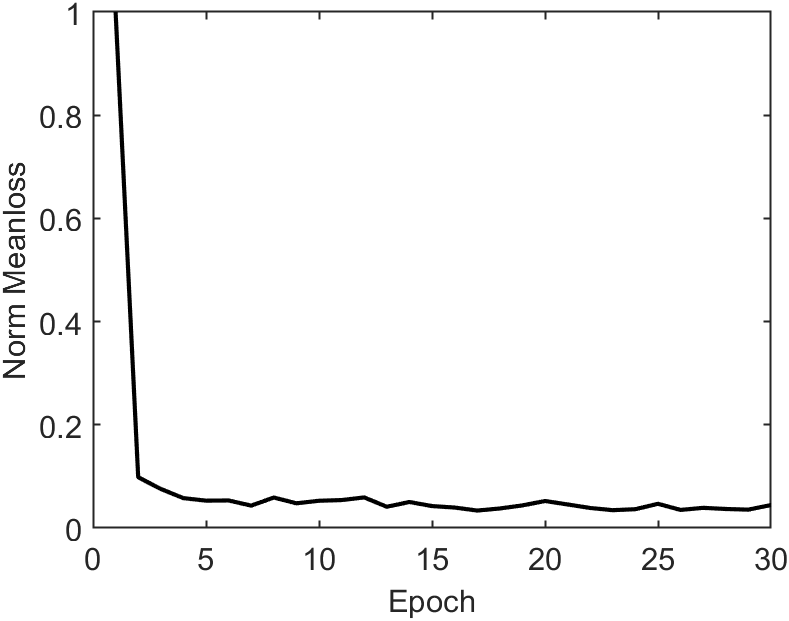

**Figure S10 Mean-loss evolution during representative live-cell calibration block training.** A typical initial fast drop within 5 epochs and thereafter stabilized mean-loss value.

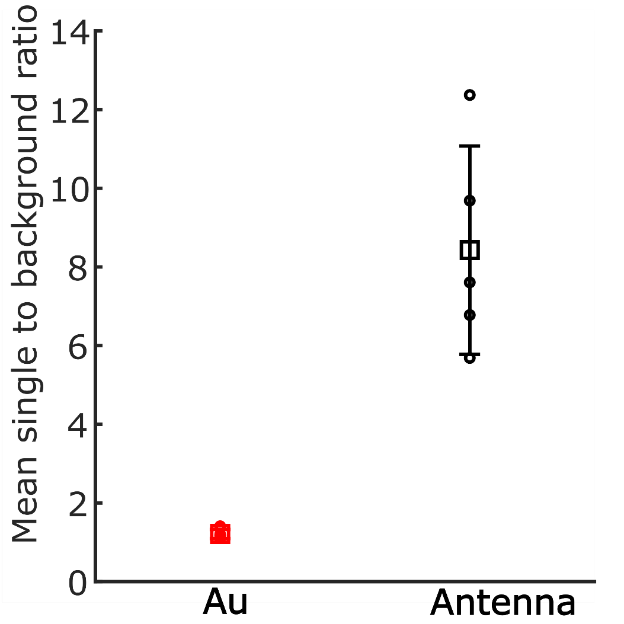

**Fig. S11** Statistical comparison of the signal-to-background ratio for the two different types of substrates. The metasurface shows a substantial increase over the control condition, demonstrating approximately one order of magnitude fluorescence enhancement relative to Au planar substrate. All fluorescence values were calculated as the average intensity within the segmented cell region.

**Note 13: Computation hardware and software**

The U-Net related training was performed in MATLAB R2024a using the Deep Learning Toolbox on a workstation equipped with an NVIDIA RTX 5000 Ada Generation GPU (32 GB memory) and an Intel Xeon w7-2495X CPU (24 cores, 48 logical processors). For the fixed-cell plasmon-guided U-Net pipeline, a representative run using 1000 raw input frames required approximately 1 min per epoch which corresponds to 3.3 h total for 200 epochs. For the live-cell block-wise calibration pipeline, each temporal block was calibrated separately and required about 30 epochs and ~4 min in total (~8 s per epoch). These runtimes show that both the fixed-cell and live-cell workflows can be executed on a single desktop workstation without requiring dedicated multi-GPU infrastructure.
